# Imputation for Lipidomics and Metabolomics (ImpLiMet): Online application for optimization and method selection for missing data imputation

**DOI:** 10.1101/2024.06.17.599353

**Authors:** Huiting Ou, Anuradha Surendra, Graeme S.V. McDowell, Emily Hashimoto-Roth, Jianguo Xia, Steffany A.L. Bennett, Miroslava Čuperlović-Culf

## Abstract

**Motivation:** Missing values are often unavoidable in modern high-throughput measurements due to various experimental or analytical reasons. Imputation, the process of replacing missing values in a dataset with estimated values, plays an important role in multivariate and machine learning analyses. Three missingness patterns have been conceptualized: missing completely at random (MCAR), missing at random (MAR), and missing not at random (MNAR). Each describes unique dependencies between the missing and observed data. As a result, the optimal imputation method for each dataset depends on the type of data, the cause of the missing data, and the nature of relationships between the missing and observed data. The challenge is to identify the optimal imputation solution for a given dataset.

**Results:** ImpLiMet: is a user-friendly UI-platform that enables users to impute missing data using eight different methods. For the user’s dataset, ImpLiMet can suggest the optimal imputation solution through a grid search-based investigation of the error rate for imputation across three missingness data simulations. The effect of imputation can be visually assessed by histogram, kurtosis and skewness analyses, as well as principal component analysis (PCA) comparing the impact of the chosen imputation method on the distribution and overall behaviour of the data.

**Availability and implementation:** ImpLiMet is freely available at https://complimet.ca/shiny/implimet/ with software accessible at https://github.com/complimet/ImpLiMet

**Supplementary information:** Supplementary data are available at Bioinformatics Advances online.

## 1 Introduction

Missing data are a major problem for multivariate, machine learning (ML) and network analyses. For example, in large lipidomic or metabolic datasets, measurements for some analytes may not be available in every sample due to routine technical variability, low abundance, ion suppression from co-eluting analytes, inaccurate feature assignment in annotation pipelines, or because analytes are simply not present in a subset of samples. This “missingness” confounds ML approaches, limits the number of methodologies that can be utilized, and reduces the statistical power of models that exclude samples with missing values. Sample exclusion further alters cohort representation, notably when “missingness” is an indicator of a particular subgroup, biasing results towards the groups in which all analytes are observed, and potentially leading to inaccurate interpretations (Jäger et al. 2021; Stoyanovich et al. 2020).

Imputing missing values is a valid solution to these problems when performing multivariate and ML analyses and can reduce data bias resulting from sample exclusion. Three types of missingness have been conceptualized that can be addressed by imputation (Mack et al. 2018; Scheffer 2002):

1. Missing completely at random (MCAR) refers to values whose absence is completely independent of any other data feature or covariate. In this type of missingness, each sample has the same probability of presenting an MCAR value because there is no underlying difference between the samples with or without missing data ((Mack et al. 2018; RUBIN 1976). A real-world example of MCAR is transient (aka random) technological failure over the course of data collection such that there is no relationship between the samples with missing or observed values.
2. Missing at random (MAR) refers to missing values whose absence is related to the values of other measured features but not to the measured values of the same feature (Schafer 1997). Here, missing values do not depend on the variable in question but on the values of the other analytes present in each sample. An example of MAR would be when the value for one analyte is missing because its measurement is obscured by the abundance of another analyte in the same sample (e.g., ion suppression of co-eluting analytes in the case of lipidomic or metabolomic datasets).
3. Missing not at random (MNAR) refers to missing values that are absent because a feature, condition, or covariate is directly responsible for the absence in that sample. Here, the probability of missingness depends on the sample itself. A biological example of this group would be analytes that are not synthesized, and thus not present, in every condition. A technological example would be when analytes are present in a given sample but are below the limit of quantification of the technology used to measure the data.

Multiple imputation methods have been introduced to approximate missing values. Recently, both Jäger et al., and Chilimoniuk et al., have compared and evaluated different approaches with respect to the quality of the imputed data and their downstream impact on ML pipelines (Chilimoniuk et al. 2024; Jäger et al. 2021). Jager et al. ((Chilimoniuk et al. 2024; Jäger et al. 2021) present a method for testing imputation quality based on the error rate and downstream use of data and in their work show that in almost all assessed examples Random-Forest (RF) provides the optimal result yet this may be not be the case for each and every dataset. To ensure agnostic dataset evaluation, Lin et al. (Lin et al. 2024) have recently presented a platform for imputation of mass spectrometry omics data that provides users with the information about the hypothetical source of missingness through correlation analysis – testing possibility for MAR and statistical analysis – and exploring the possibly for missing through MNAR mechanisms. Users can then provide the ratio of missingness types present in their datasets, that will influence the selection of the imputation method, however the same imputation method is used for all variables. A remaining bioinformatic challenge is the identification of the optimal imputation solution for a given dataset of any type. As missingness can come from different sources for variables within the dataset, different imputation methods might be necessary for groups of features within the dataset.

To address this challenge, we present **Imp**utation for **Li**pidomics and **Met**abolomics – ImpLiMet – applicable to any numerical data set, validated here for using lipidomic and metabolomic data. ImpLiMet is an R package available at https://github.com/complimet/ImpLiMet and online with UI at Computational Lipidomics and Metabolomics: CompLiMet: https://complimet.ca/shiny/implimet/. ImpLiMet enables users to impute missing data using eight different methods across the whole dataset or within user-defined groups of features. The effect of each method can be visualized by histogram, kurtosis and skewness analyses, as well as principal component analysis (PCA) comparing the impact of simply removing features and samples with missing data to the chosen imputation method. To identify the optimal imputation solution, ImpLiMet further offers an optimization option wherein the error of each imputation method is evaluated, and the user is informed of the method with the lowest mean absolute percentage error (MAPE) across three “missingness” simulations for their dataset.

## 2 Implementation

ImpLiMet is written in R and deployed with a RShiny UI. It is compatible with all modern web browsers. Figure 1 presents the ImpLiMet workflow and pseudocode for the optimization procedure. To run ImpLiMet, the user uploads their dataset as a .CSV file. If the dataset includes features measured in different units by different platforms (multiple feature measurement groups) or features possibly having different levels of relationships to other features, the user has the option to format their data such that the imputation methods consider feature groups separately. An example of different measurement groups could be the combination of lipidomic and metabolomic data measured using different platforms or multiomics data with, for example. metabolomic and transcriptomic data contained in a single dataset. The user can then specify the number of features or samples with the selected percentage(s) % of missing values to be removed prior to choosing an imputation measure or optimizing across measures. Eight imputation methods are available: 1) replacing with the feature minimum, 2) replacing with the feature minimum divided by 5, 3) replacing with the feature maximum, 4) replacing with the feature median, 5) replacing with the feature mean, 6) using K-Nearest Neighbors (kNN) (Hastie, et al. 1999; Troyanskaya, et al. 2001), 7) using Random Forest (RF) (Pantanowitz, Marwala, 2009) or 8) using Multivariate Imputation by Chained Equations (MICE) (van Buuren et al. 2011). For kNN, RF, and MICE, the user is queried to provide the number of neighbours for kNN, the number of trees for RF, and for MICE the number of iterations. kNN is implemented using *impute*.*KNN* function; RF imputation utilizes *missRanger*.*RF* function (Stekhoven, Buehlman, 2011) and MICE using the function *mice* (van Buuren et al. 2011).

**Figure 1.**
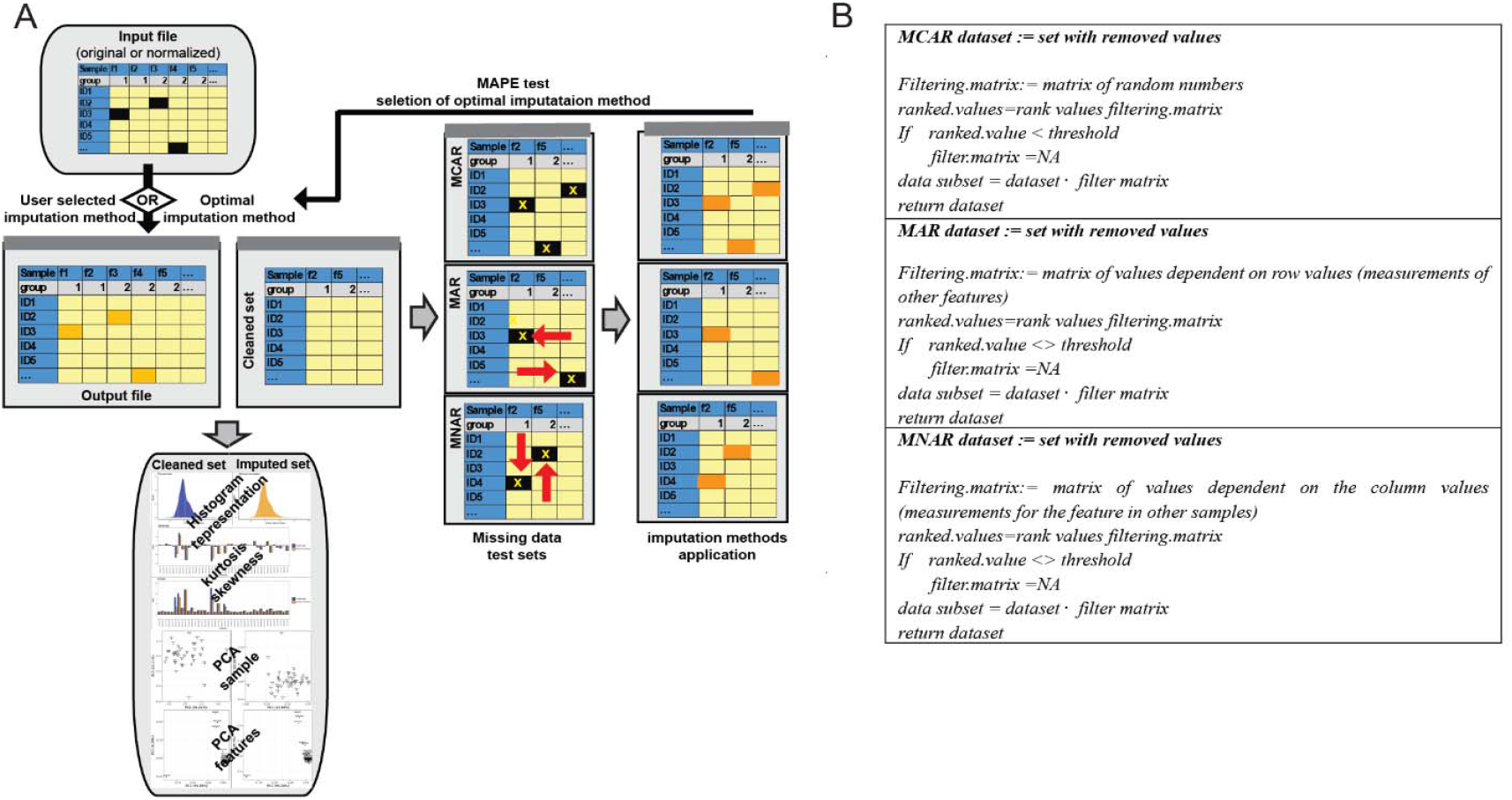
A. Schematic workflow of ImpLiMet. In the case of automated optimization, ImpLiMet first removes all columns in the dataset with missing values then simulates missing elements following three types of missingness: MCAR, MAR and MNAR. Missing values are imputed with all methods and the error of imputation is determined using MAPE. Imputation is then performed on the original dataset using the method with the lowest MAPE value. The dataset with imputed values is returned to the user and the effect of imputation on the dataset is visualized with statistical measures and PCA. B. Schematic pseudo code of the process of data removal for the three different missingness types during optimization. Matrix multiplication indicates the element-wise product. Detailed pseudo code is provided in the Supplementary Materials. A comprehensive flowchart is presented at: https://complimet.ca/shiny/implimet/

If the user’s dataset has at least 3 features and 6 samples with no missing values, or a minimum of 18 non-missing values across minimum of 3 features and 6 samples, ImpLiMet further offers an optimization option wherein the error of each imputation method is evaluated by simulating the three different sources of missingness in the user’s dataset once all missing data is removed then testing all available imputation methods. Optimization suggests, as the best imputation method, the one with the lowest mean absolute percentage error (MAPE) across the three “missingness” data simulations, i.e., the lowest value for all tested values. The selected approach is used to impute the original dataset and this result is provided as a download. Alternatively, the user can choose to utilize another imputation method based on, for example, simulation results, the visualization analysis provided by ImpLiMet, or prior information about the sources of missingness in the dataset. In the case of different types of missingness in the dataset, the user can group features by missingness type, perform imputation using the proposed optimal methods for each group and subsequently combining the results for different groups using the downloaded data.

In the optimization step, samples without any missing values are selected to create a complete set. If the cleaned dataset obtained by removing all samples (rows) with missing values has no remaining values, optimization instead selects features (columns) without missing values. Finally, if both approaches result in the removal of all columns and rows, ImpLiMet selects columns and features with less than 80% missing values and, with the remaining set, returns to selecting samples with no missing values. If not found, then ImpLiMet selects features with no missing values. In this way. the algorithm ensures that the analysis of the optimal imputation method for the dataset can be evaluated by imputing only the missing data from the set that is removed for testing in the optimization step. Note, if there are less than at least 18 values, in 6 samples and 3 features remaining, optimization of imputation cannot be done. It is important to keep in mind that in extremely small datasets imputation will be biased by available information. From the dataset devoid of missing features, ImpLiMet removes data values at the sample threshold percentage initially provided by the user for filtering. If threshold percentage is not provided, i.e., user opts not to remove any additional features or samples from their dataset prior to imputation, ImpLiMet uses 30% as the threshold percentage in the optimization process. The threshold percentage is used to simulate the optimal imputation method for a given dataset at the level of the user’s specified tolerance for imputation. For extremely small dataset sizes (e.g., a 6×3 matrix), only a 10% threshold for full optimization will enable simulation as all other thresholds will result in an insufficient sample size for imputation method testing and error calculation. The known values removed for simulation are kept as the hold-out set and are used to evaluate error of imputation as follows:

Given dataset: 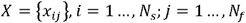 where *N*_*s*_ is the number of samples and *N*_*f*_ is the number of features; with missing elements *x*_*km*_, (*k,m*) ∈ *M* the goal of imputation is to determine values for the missing elements that resemble the complete data. As the first step in optimization, any row or column with missing elements are removed leading to the subset 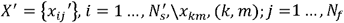.

From this subset data, removal is performed separately to simulate MAR, MCAR, and MNAR mechanisms. Pseudocode for each missingness mechanism is provided in Figure 1B.

For **MCAR**, a filtering matrix of dimension *N*′_*s*_ x*N*′_*f*_ is created by random sampling from a uniform distribution (minimum = 0 and maximum = 1) generated from the function *runif* in R. Random values in the matrix are ranked and values below the imputation threshold are set to NA for missing and above are set to one for remaining. The element-wise product between this filtering matrix and full data matrix provides the MCAR example set for further testing.

For **MNAR**, the missing value assignment is performed individually for each feature as follows: 1) A list of values is generated by sampling from a logistic distribution (location = 0, scale = 1), denoted 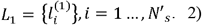 A second list is generated by sampling from the uniform distribution (minimum = 0 and maximum = 1), denoted 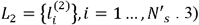 A third list is generated from the product of 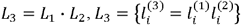. 4) The ranks for the values in *L*_1_, *L*_3_, as well as the feature measurements, are computed. 5) The highest and lowest ranks from *L*_3_, with the number of missing values dependent on the assigned threshold, are determined and the corresponding (feature-wise) ranks in *L*_1_ are assigned. Equivalent ranks in the data set are removed as missing.

For **MAR**, a co-dependance group is created by summating all feature values in a sample except the values in the current cell. If the input file contains information about the feature groups, based on biological or analytical characteristics, the summation calculation is performed within each feature group for each sample for the co-dependence matrix. The MAR process follows MNAR steps 1 through 3. In step 4, the ranks for the values in *L*_1_, *L*_3_, and the sample values in the filtering matrix are computed. Missing indices are assigned to the highest and lowest ranks from *L*_3_, with the number of missing values dependent on the sample threshold. The order of the values in *L*_1_, that produces the missing indices in *L*_3_, are retrieved, and the corresponding order in the filtering matrix column for the co-dependent feature are assigned as NA.

After generating the three types of missing datasets, each dataset is imputed using each of the eight available methods. For multivariate methods, users are prompted to select a simple or full version of parameter optimization. Simple parameter optimization uses the following default parameters: K-value=10, Tree Value=500, and Mice Iteration=2. If a full parameter search is selected, the accuracy of the imputed values is tested over a range of hyperparameters for kNN, MICE and RF. Specifically, for kNN, the K-values tested range from 10 to 100 incremented by 20. For the optimal K-value in in this range, a refined search is conducted from k-4 to k+4 in single value increments to identify the K-value with the lowest error rate. For RF, the number of trees in the sequence of 5, 10, 20, 50, 100, 150, 200, 500 are examined to determine the optimal tree size. For MICE, 1-3 iterations are tested. The full optimization approach is generally preferred, however due to the large number of calculations taken in this approach it can be time consuming (e.g., for dataset with 45 samples x 40 features – the example input set provided – full optimization test takes ∼2 minutes online). Thus, for very large datasets, fast optimization analysis can provide initial screen of methodologies. Error rates are calculated by mean absolute error rate (MAPE) defined as:

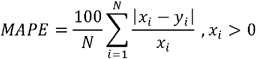

where *N* is the number of missing values, *x*_i_ is the actual value and y_i_ is the prediction. The MAPE results for each of the eight imputation methods assessed for each missingness mechanism are displayed and the method with the lowest MAPE value across the missingness mechanisms is highlighted and used for imputation. In general, omics measurements are greater than zero as the minimal value measured corresponds to the minimal level of detection in the measurement, rather than absolute zero value.

The effect of imputation on the dataset is visualized by dataset histogram, kurtosis and skewness characteristics as well as PCA comparing the original dataset, following removal of all samples and features with missing data, to that of the imputed dataset. Histograms show all values in the dataset following feature z-score scaling and compares the overall dataset distribution of cleaned dataset with the imputed set. Kurtosis and skewness provide information about the distribution for each feature separately. Kurtosis is a measure the level of tailing of the data. Skewness indicates the symmetry relative to the normal distribution. Symmetric data has a skewness of zero. High negative skewness indicates that data are left skewed (a long-left tail, thus data are missing more values in the high abundance range). Positive skewness indicates data are right-skewed, meaning that more low abundance data are missing altering the assumption of a normal distribution. High skewness, calculated in ImpLiMet using R function *skewness*, suggests the possibility of MNAR for those features. Kurtosis (calculated using R function *kurtosis*), indicates potential increased levels of outliers in the dataset, with high values suggesting significant presence of outliers from normal distribution. In ImpLiMet, kurtosis and skewness are shown for both datasets with all samples and features with missing values removed and the complete, imputed dataset, allowing the user to explore possibility for of MNAR in some of the features as well as to observe the effect of imputation on the normality of features distribution. PCA, for both samples, calculates principal components using features as variables, and displays features, using their values across samples as variables. The user-provided sample and feature names are shown in the plots for reference. An example of the optimization utilization as well as comparison of errors in imputation using recommended and other imputation methods is presented in the Supplementary Materials.

Briefly, from the subset of metabolomics data published by Li et al. (Li et al. 2019) with complete data for 50 samples and 50 features, we have removed values from 120 cells and tested the error rate for the imputed values using different methods. Results show that the recommended method, in this case RF, provides imputation with the lowest error and the best agreement in PCA when comparing the original dataset with the original data (full information is provided in Supplementary Materials). Furthermore, in the supplementary information we provide an example of the utilization of ImpLiMet on a combined metabolomics and lipoprotein dataset (Oppong et al. 2024) with multiple groups and testing of the skewness analysis.

## 3 Conclusion

ImpLiMet is a versatile and open-access Web-based application designed to assist users in identifying the optimal imputation solution for their given dataset. The imputation recommendation returned to the user shows the optimal method based on the lowest error rate overall, while at the same time presenting error rates of imputation for different types of missingness for all methods. ImpLiMet currently includes eight previously presented imputation methods as well as visual representation of statistical features of the dataset helping users interpret sources of missingness across features. ImpLiMet provides a possibility for distinct feature groups and includes a visual assessment of data properties related to the missingness type in addition to the optimization of imputation analysis. Future interations will include the addition of other imputation methods as well as an automated analysis of the type of missingness present in the dataset and automated multi-method imputation across groups.

