## Supplementary Materials for "Imputation for Lipidomics and Metabolomics (ImpLiMet): Online application for optimization and method selection for missing data imputation"

### S1. Optimization method performance testing

Imputation methods included in ImpLiMet have all been previously developed, tested, and extensively used (Chilimoniuk, et al. 2024; Hastie et al. 2000; Pantanowitz and Marwala, 2009; Stekhoven et al. 2011; Troyanskaya et al. 2001; van Buuren et al. 1999; van Buuren et al. 2006; van Buuren et al. 2011; Wright and Ziegler, 2017). If *Optimization* is selected in the analysis, ImpLiMet determines the imputation error rate for different methods and suggests to the user the best performing imputation method for the dataset. The optimal method for imputation for a given dataset is performed through a grid search across all methods and with range of hyperparameters. The error level is determined for three different types of missingness: Missing completely at random (MCAR), Missing not at random (MNAR), and Missing at random (MAR). Hyperparameter values used in the optimization search are shown in Supplementary Table 1.

**Supplementary Table 1.** **Hyperparameter values included in the optimization of machine learning imputation methods.**

| Method | KNN | RF | MICE |
| --- | --- | --- | --- |
| Type of Parameter | K-value | Tree value | Number of iterations |
| Optimization:  Full search | 10:100 (20)  Fine search: Min-4:Min+4 (1) | 5, 10, 20, 50:200 (50), 500 | 1:3 (1) |
| Optimization:  Fast search | 10 | 500 | 2 |

The range of values is shown. The step value for the list is indicated in brackets.

Imputation error is calculated by mean absolute error rate (MAPE) defined as:

$$MAPE=\frac{100}{N}\sum_{i=1}^{N} \frac{\left| x_{i}-y_{i} \right|}{x_{i}}$$

where *N* is the number of missing values, $x_{i}$ is the actual value and $y_{i}$ is the prediction.

Imputation optimization is done using a subset of data that does not have any missing values. In this case ImpLiMet first removes samples (rows) with any missing values. If the resulting subset has less than 6 rows (samples), ImpLiMet instead removes all features (columns) with any missing values from the original dataset. If the remaining set has less than 3 features and less than 6 samples, the optimization step cannot be performed. In this case, the user can still select their preferred imputation method and perform imputation on the original set. If 6 or more samples and 3 or more features in the provided set are complete, i.e., they have no missing values, then these are selected and used for the imputation method error testing and optimization. To determine the error rate, missing data are simulated by removing cells from the cleaned dataset (the previously selected samples and features without missing data) following the procedures described in Supplementary Box 1.

**Supplementary Box 1.** Overview, pseudo code showing methods for cell removal that represent different missingness types. All matrix products correspond to element-wise matrix products.

| ***MCAR dataset := set with removed values***    *Filtering.matrix:= matrix of random numbers*  *ranked.values=rank values filtering.matrix*  *If ranked.value < threshold*  *filter.matrix =NA*  *data subset = dataset* $\cdot$ *filter matrix*  *return dataset* | ***MNAR dataset := set with removed values***    *For i=1:number of columns*  *L1:= logistic distribution random set*  *L2:=uniform distribution random set*  *L3 = L1* $\cdot$ *L2*  *ranked.L3 (1:row,i)=rank values L3*  *ranked.L1(1:row,i)=rank values L1*  *ranked.values(1:row,i)=*  *rank column values*  *missing.rank=*  *ranked.L1*$\hat{=}$*ranked.L3<>threshold*  *filter.column=NA, when*  *ranked.values eq missing.rank*  *data subset = dataset* $\cdot$ *filter matrix*  *EndFor*  *return dataset* | ***MAR dataset := set with removed values***  *For i=1:number of columns*  *L1:= logistic distribution random set*  *L2:=uniform distribution random set*  *L3 = L1* $\cdot$ *L2*  $sum.row=\sum_{\forall row \backslash\left\{ current \right\}} values$  *ranked.L3 (1:row,i)=rank values L3*  *ranked.L1(1:row,i)=rank values L1*  *ranked.values(1:row,i)=*  *rank sum.row*  *missing.rank=*  *ranked.L1*$\hat{=}$*ranked.L3<>threshold*  *filter.column=NA, when*  *ranked.values eq missing.rank*  *data subset = dataset* $\cdot$ *filter matrix*  *EndFor*  *return dataset* |
| --- | --- | --- |

Missing data is imputed using all methods included in ImpLiMet. The difference between the imputed and the original values in the cleaned set is calculated using the MAPE formula. The minimal MAPE value is suggested as the optimal method and is used to impute the dataset’s existing missing values. A table of MAPE values for the three different missingness approaches and all imputation methods is provided. The imputation recommendation depends on the characteristics of samples as well as type of missingness and the sample size. The optimization method is a simple grid search identifying the method that provides the lowest error rate across all missingness patterns in their specific dataset.

An example of the performance of the imputation method optimization is shown using metabolomics dataset published by Li et al. (Li et al. 2019). The subset used in this example measures 50 samples and 50 features. The selected set does not have any missing values. PCA for the complete dataset is shown in Supplementary Figure 1A. From this dataset we have removed values from 120 cells, PCA for the cleaned dataset, the subset of values with no missing value selected following random deletion of 120 values is shown in Supplementary Figure 1B. On this dataset we ran optimization and imputation with the recommended method as well as all other methods. Imputation results for different methods are compared using MAPE calculation for the imputed and original values, prior to removal, as well as comparison results using PCA results. Results are shown in the Supplementary Figure 1.

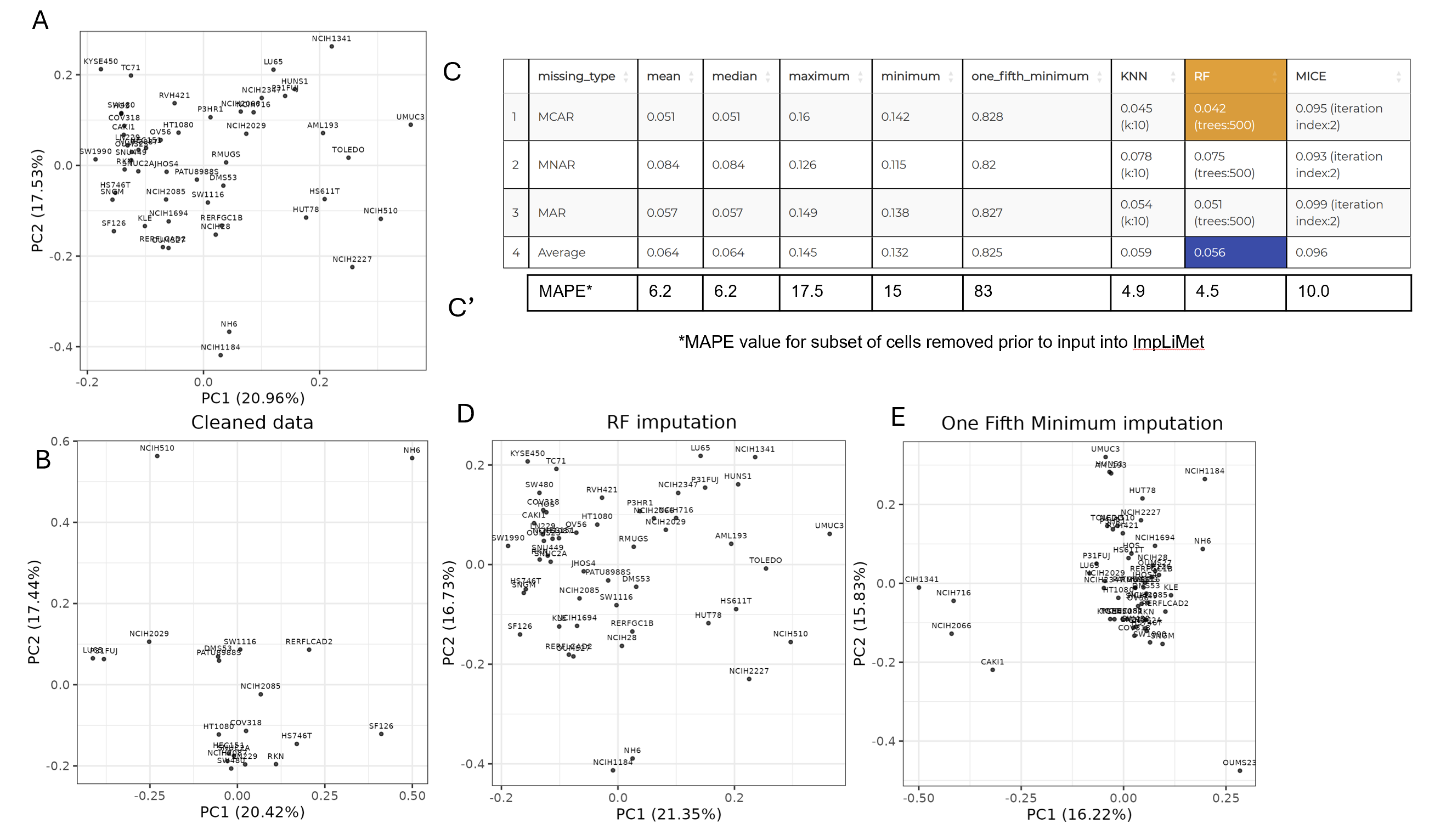
**Supplementary Figure 1.** Example of optimization and imputation presented on a subset of the metabolomics data of (Li et al. 2019). Subset includes 50 samples and 50 features with no missing values. A. PCA for the complete, original set used in the analysis. From this set we randomly removed 120 values across different samples and features for imputation analysis. B. PCA of the subset with only complete rows and columns after the removal of 120 values from the set shown in A. C. For this dataset ImpLiMet optimization recommends Random Forest (RF) as the optimal imputation method as this approach leads to the lowest MAPE value overall in MCAR missingness type. C’ As in this case we have the original values for the 120 cells removed in the analysis, we are also including here MAPE values for the imputed and original values, prior to removal. D. PCA of the optimized imputation result using RF, the lowest MAPE value method. E. PCA of samples following the imputation using 1/5^th^ of the minimum value which is in this case has the highest MAPE values.

This benchmark analysis shows that error in imputation values for the missing data fully agrees with the MAPE values obtained in the optimization process. The recommended method, in this case Random Forest (RF) has the lowest error rate based on MAPE analysis using imputed values and values in the original dataset. Furthermore, PCA of the RF imputed dataset is in the agreement with the original dataset’s PCA – with highly comparable PC1 and PC2 values (Supplementary Figure 1. A and D) while PCA values following 1/5^th^ of the minimum value imputation, as a suboptimal method in this dataset, show different PC1 and PC2 values and PCA projection structure (Supplementary Figure 1. E).

### S2. Example: Metabolomics dataset imputation

In this example we show utilization of Implimet for the imputation of a dataset previously published by Oppong et al. (Oppong, 2024). Briefly, this dataset contains metabolomics and lipoprotein measurements in serum of 191 patients with relapsing-remitting Multiple Sclerosis (MS) (RRMS, N=52), patients with neuromyelitis optica (DCs, N=30), secondary progressive MS (SPMS, N=29), and control healthy donors (HC, N=80). Included in the analysis are both metabolites and lipoproteins measured in serum using a high throughput NMR spectroscopy platform. Study design and detailed analysis are provided in the original publication. This dataset had several values set to zero, and thus, prior to imputation testing all zero values in the dataset are set to missing values. Additionally, for this application presentation here for three features we have respectively removed data that are below a threshold value, above a threshold or randomly selected through the feature set. Specific features are shown in Supplementary Figure 2. A set of 75 metabolites and lipoproteins was selected for this demonstration with metabolites and lipoproteins in this dataset separated into two groups such that imputation was performed only within a group of features (adding as a second row named “groups” with group labels). Although in this set both groups of features are measured using NMR methodology, they required two different pulse sequences for analysis of small molecules (metabolites) and large constructs (lipoproteins). Following the upload ImpLiMet shows dataset to have 191 samples, 75 features and 296 missing values. In this analysis we are selecting to not remove any samples and features prior to imputation. Optimization investigation in this dataset indicates that imputation using RF with 5 trees has the lowest MAPE value overall obtained for the MCAR missingness type. Optimization table is shown in Supplementary Figure 2A. We have selected full optimization which is using hyperparameters listed in Table 1.

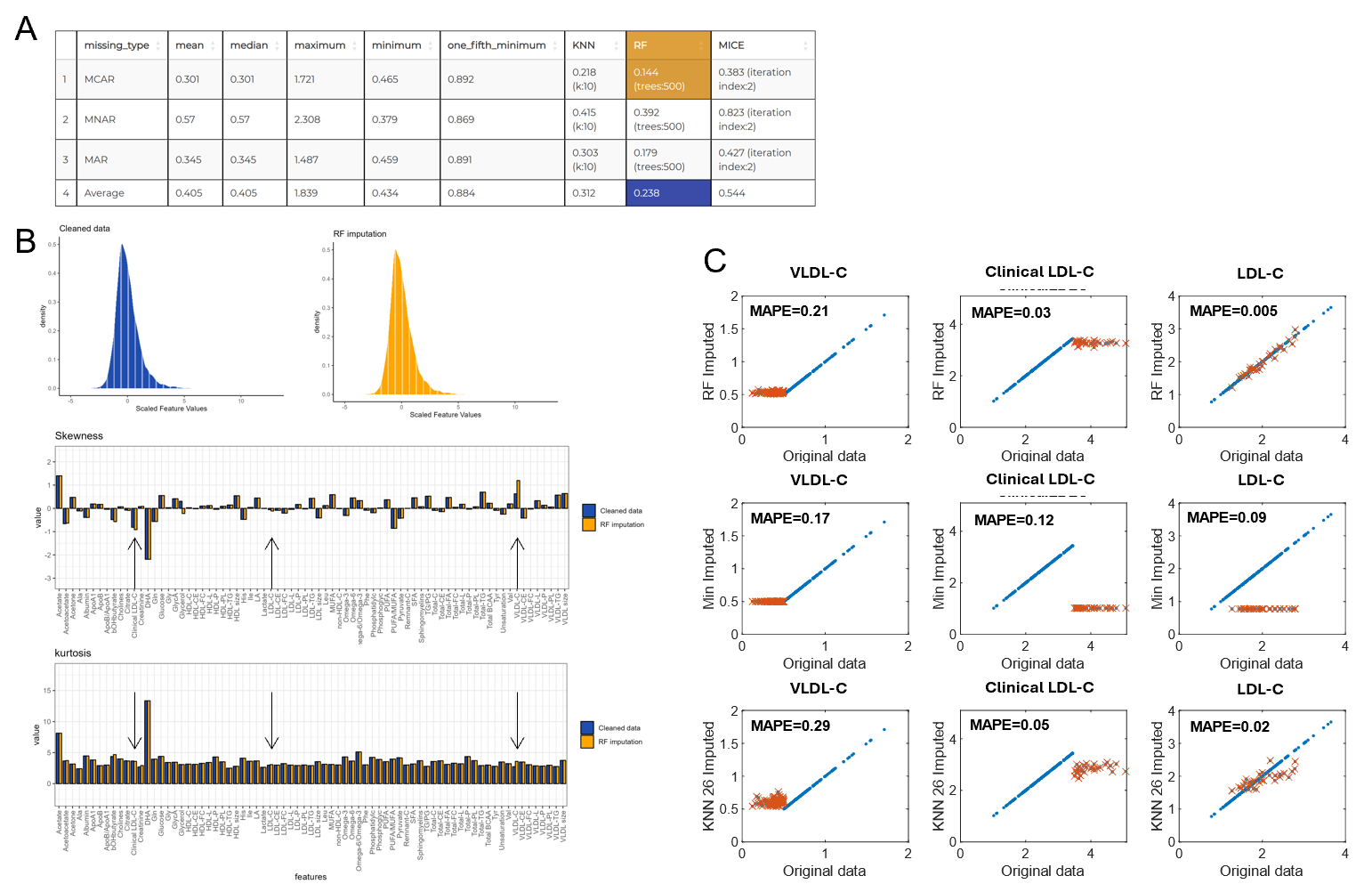

**Supplementary Figure 2.** Imputation analysis using ImpLiMet presented on the Multiple Sclerosis dataset published by (Oppong, 2024). A. A full parameter search analysis of the optimal method for imputation on this dataset. Indicated and used for subsequent imputation is the method that has the lowest MAPE value in all types of missingness, in this case RF with 5 trees. B. Visual outputs of some statistical characteristics of the input dataset with features with missing values removed (Cleaned data) and dataset following RF imputation. Shown are data values histograms, skewness and kurtosis analysis for each feature separately. C. For method presentation in this example we have deliberately removed values below 0.5 in VLDL-C measurements, above 3.5 in Clinical LDL-C and randomly 30 values of 191 for LDL-C. Plots compare original values for these three features with values obtained in imputation (red) as well as all the other values (blue, unaffected by imputation). Shown are results for RF Imputation, min value imputation and KNN with 26 neighbours. Each plot shows MAPE values for the feature.

For larger dataset or faster screening, it is possible to do optimization on a single hyperparameter (without selecting the full parameter search in the options for optimization) but whenever possible it is recommended to do the full test. Supplementary Figure 2B shows some of the result visualization provided by ImpLiMet. Histogram for the cleaned data where features with missing values are removed compared to the set with RF Imputed data shows no change in the overall value distribution. As there are only 296 values missing out of total of 14325 values, this is expected. Skewness and kurtosis values are shown for each feature separately for the two datasets (Supplementary Figure 2B). For VLDL-C and Clinical LDL-C, where values are removed prior to imputation for values below or above a threshold, there is a slight increase in absolute value of skewness following imputation. Comparison of original values and values obtained with different types of imputation (Supplementary Figure 2C) clearly shows reasons for this increase in the distribution skewness where imputation leads to values around the threshold. The RF imputation shows for these specific features minimal MAPE values, with particularly low error rate in the example of LDL-C, where values have been removed completely at random. For the VLDL-C, values below concentration of 0.5 are removed, RF imputation result matches minimal value in the remaining set. For Clinical LDL-C, where values over 3.5 have been removed, RF values are largely matching the maximum remaining value, clearly leading to an increased in skewness. Thus, for VLDL-C, imputation with minimum value leads to a highly comparable MAPE value with RF imputation. Thus, although optimization analysis provided by ImpLiMet does not test for the missingness source in the dataset, analysis of skewness and kurtosis can be used by user to determine possible sources of missingness through the investigation of the left or right-side skewness of the original and imputed dataset.

### S3. ImpLiMet web application

(a) **Input File format**

**Table 2. Example of the ImpLiMet input file format required if the dataset has only one feature measurement group**

| Sample | feature1 | feature2 | feature3 | feature4 | … |
| --- | --- | --- | --- | --- | --- |
| ID1 | 15669.4 | 205.2 | 56.5 | 361.5 | 12 |
| ID2 | 10084.3 | 220.9 |  | 438 | 8.3 |
| ID3 | 12836.6 | 394.7 | 93.9 | 861.2 |  |
| … | 10520.2 | 293 | 200.1 | 1309.9 |  |

Row 1 must contain feature names. Column 1 must contain unique sample IDs. Missing values should be indicated as NA or as empty cells.

**Table 3. Example of the ImpLiMet input file format required if the user includes information about multiple feature measurement groups**

| Sample | feature1 | feature2 | feature3 | feature4 | … |
| --- | --- | --- | --- | --- | --- |
| group | 1 | 1 | 1 | 2 | 2 |
| ID1 | 15669.4 | 205.2 | 56.5 | 361.5 | 12 |
| ID2 | 10084.3 | 220.9 |  | 438 | 8.3 |
| ID3 | 12836.6 | 394.7 | 93.9 | 861.2 |  |
| … | 10520.2 | 293 | 200.1 | 1309.9 |  |

If the dataset includes features measured in different units by different platforms (multiple feature measurement groups), data should be formatted to indicate which groups must be considered separately for missing data simulation (i.e., which data were measured in the same units on the same platform). In this case Row 1 must contain feature names. Row 2 must contain the group information. Column 1 must contain the sample IDs. Missing values should be indicated as NA or as empty cells.

(b) **Running ImpLiMet**

The user uploads the dataset, selects the percentage threshold for imputation. After threshold is selected, the user can select type of imputation method that will be used. If the optimization option is selected, user can further select the full parameter search. With this option the calculation of MAPE value for three types of missingness is performed using hyperparameters listed in Table 1.

(c) **Output**

The ImpLiMet output includes the imputed dataset as well as a visualization of the effect of the chosen (or optimized) imputation method on the dataset. The visualization tabs provide histograms for cleaned and imputed datasets as well as comparison of kurtosis and skewness values for each feature in the original and imputed datasets. For visualization of overall effect of imputation ImpLiMet also shows PCA plots of the cleaned dataset (i.e., dataset with all columns and rows with any missing value removed) and of dataset imputed with the selected method. PCA is performed on both samples and features with the names of samples and features included in the plot for easy reference. The optimization result is shown in the MAPE table. The method with the lowest MAPE for the dataset across three missingness types is highlighted and used for imputation.
